# Both mOTS-words and pOTS-words prefer emoji stimuli over text stimuli during a reading task

**DOI:** 10.1101/2023.11.07.565794

**Authors:** Alexia Dalski, Holly Kular, Julia G. Jorgensen, Kalanit Grill-Spector, Mareike Grotheer

**Affiliations:** Department of Psychology, Philipps-Universität Marburg, Marburg 35039, Germany; Center for Mind, Brain and Behavior – CMBB, Philipps-Universität Marburg and Justus-Liebig-Universität Giessen, Marburg 35032, Germany; Department of Psychology, Stanford University, Stanford, CA 94305, USA; Wu Tsai Neurosciences Institute, Stanford University, CA 94305, USA

**Author notes:** **Corresponding author:** Alexia Dalski.

## Abstract

The visual word form area in the occipitotemporal sulcus (OTS), here referred to as OTS-words, responds more strongly to text than other visual stimuli and is crucial for reading. We hypothesized, that this text preference may be driven by a preference for reading tasks, as in most prior fMRI studies only the text stimuli were readable. Hence, we performed three fMRI experiments (N=15) and systematically varied the participant ‘s task and the stimulus, investigating mOTS-words and pOTS-words subregions. In experiment 1, we contrasted text stimuli with non-readable visual stimuli (faces, limbs, houses, objects). Experiment 2 utilized an fMRI adaptation paradigm, presenting compound words in text or emoji formats. In experiment 3, participants performed a reading or a color task on compound words in text or emoji format. Using experiment 1 data, we identified mOTS-words and pOTS-words by contrasting texts with non-readable stimuli. In experiment 2, pOTS-words, but not mOTS-words, showed fMRI adaptation for compound words in both text and emoji formats. In experiment 3, surprisingly, both subregions showed higher responses to compound words in emoji than text format. Moreover, mOTS-words showed higher responses during the reading than the color task and a task-stimulus interaction. Multivariate analyses revealed that distributed responses in pOTS-words encode the visual stimulus, while responses in mOTS-words encode both stimulus and task. Together, our findings suggest that the function of the OTS-words subregions goes beyond the specific visual processing of text and that these regions are flexibly recruited whenever semantic meaning needs to be assigned to visual input.

**Significance Statement:** Reading skills are essential in modern society and supported by a brain region in the occipitotemporal sulcus (OTS-words) that is critical for fluent reading. Here we evaluated if responses in OTS-words are driven by the activity of reading or simply by seeing text or other readable symbols (emojis). We found that OTS-words can be divided into two sub-regions which both prefer readable emojis over text, whereas the anterior sub-region also prefers reading over other tasks. These results suggest that OTS-words is flexibly recruited to encode any readable visual stimulus, not just text. By demonstrating OTS-words ‘ flexibility, this work reconciles previous contradictory findings on this regions ‘ functional properties and inspires future research on OTS-words, including its emergence during literacy acquisition.

## Introduction

Reading text is a fundamental building block of communication in our society and an inability to do so has profound negative consequences (Weiss et al., 1991; Cho et al., 2008; Ardila et al., 2010). An essential neural substrate of reading is the visual word form area (VWFA; Cohen et al., 2000), a left-lateralized region (Cai et al., 2008; Dehaene et al., 2010; Strother et al., 2017; Canário et al., 2020) which responds more strongly to text than other visual stimuli (Cohen et al., 2000; Dehaene & Cohen, 2011) and is critical for reading (Gaillard et al., 2006; van der Mark et al., 2009; Sandak et al., 2018). Here we refer to this region as OTS-words due it its anatomical location in the occipitotemporal sulcus (OTS) and its functional preference for words.

Importantly, the functional role of OTS-words is not without debate. While the predominant view suggests that it is selectively involved in processing text (Cohen et al., 2000; Cohen & Dehaene, 2004; Szwed et al., 2011), several studies found evidence against an exclusive text preference (Ben-Shachar et al., 2007; Mei et al., 2010; Vogel et al., 2012; Wright et al., 2008; for reviews see Price & Devlin, 2003; Vogel et al., 2014). Recent work further proposes a posterior-to-anterior increase in processing level along the OTS (Caffarra et al., 2021; Carreiras et al., 2014; Thesen et al., 2012; Vinckier et al., 2007), and in fact, it has been suggested that OTS-words can be divided into two subregions (middle mOTS-words and posterior pOTS-words) that differ in their functional properties, white matter connections, and functional connectivity (Stigliani et al., 2015; Lerma-Usabiaga et al., 2018; Yablonski et al., 2023). Here we hypothesize that the debate on the functional role of OTS-words may be resolved by evaluating the two sub-regions separately, as it is possible that only one of the subregions is selective for text. Moreover, we postulate that it is important to also take the participants ‘ task into consideration. In many prior experiments only the text stimuli were readable, so the observed text preference may have been confounded with a preference for reading tasks. Indeed, recent studies have found higher responses for reading than other tasks in OTS-words supporting this notion (Grotheer et al., 2018; Qu et al., 2022; White et al., 2023). We hence derived three hypotheses (Fig. 1a-c): H1: Both OTS-words subregions prefer text over other stimuli, irrespective of task. H2: Both OTS-words subregions prefer reading over other tasks, irrespective of the stimulus. H3: There is an anterior-to-posterior gradient in processing level, whereas mOTS-words is selective for reading and pOTS-words is selective for text stimuli.

**Figure 1.**
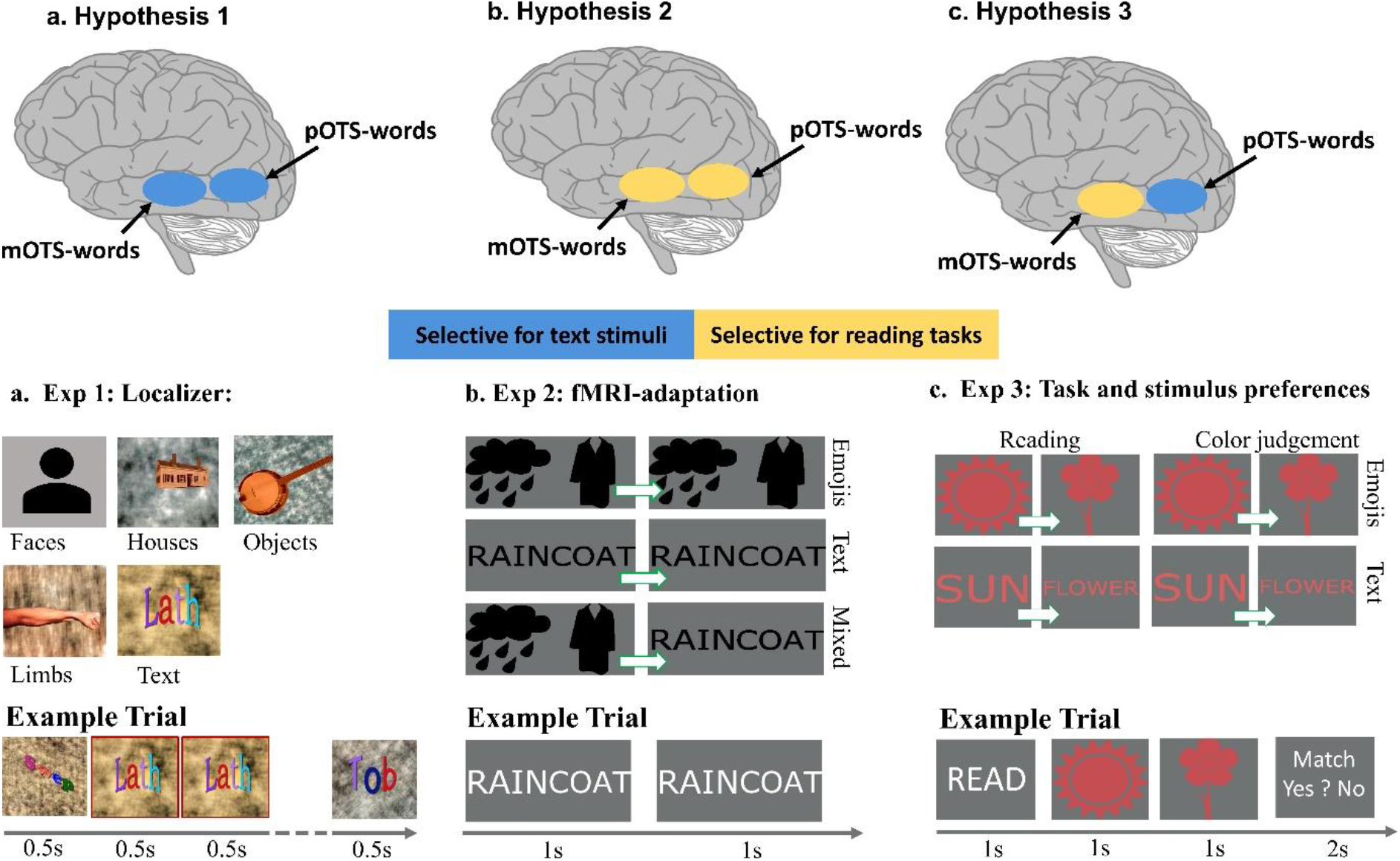
Summary of hypotheses and experimental design. **a. Hypothesis 1**. Both subregions of OTS-words show a preference for text stimuli. **b. Hypothesis 2**. Both subregions of OTS-words prefer reading tasks over other tasks. **c. Hypothesis 3**. pOTS-words is selective for the visual features of text and mOTS-words is selective for reading tasks. **d. Exp 1: Localizer**. Five visual categories (faces, houses, objects, limbs, text) were presented while subjects performed a 1-back task. Images were either colored or in grayscale. Face stimuli in the experiment were photographs of faces presented on the same scrambled background. **e. Exp 2: fMRI-adaptation**. Two pairs of images were presented, whereas each pair formed an English compound word; Task: Participants had to indicate when the images were white rather than black. All images in a trial were either in emoji, text, or mixed format. Pairs in a trial could be repeated or non-repeated. **f. Exp 3: Task and stimulus preferences**. Subjects were presented with two stimuli, presented one after the other, in emoji or text format. Tasks: Subjects were instructed to either read the word pair and indicate if they form a meaningful English compound word (reading task) or to compare the color hues of the two stimuli (color task).

To distinguish between these hypotheses, in three fMRI experiments, we systematically varied the stimulus and the participant ‘s task and assessed mean and distributed responses in mOTS-words and pOTS-words (Fig. 1d-f). In experiment 1, we contrasted responses to text with other visual categories. We used experiment 1 data to identify mOTS-words and pOTS-words based on their text preference, as in previous studies (Stigliani et al., 2015; Rosenke et al., 2021). In experiment 2, we used fMRI adaptation to probe the stimulus sensitivity of mOTS-words and pOTS-words, comparing text and emoji stimuli. In experiment 3, subjects performed a reading and a color task on readable text and emoji stimuli.

Our data show that mOTS-words and pOTS-words can be identified based on their preference for text. Strikingly, however, even though regions were identified through their sensitivity towards text, our univariate analyses showed higher responses for emojis than texts in both subregions. In pOTS-words we further observed fMRI adaptation for both text and emoji stimuli, whereas mOTS-words showed a preference for reading over a color task. Multivariate analyses align and showed that pOTS-words encodes the visual stimulus, while mOTS-words encodes both the visual stimulus and the performed task. These results expand our understanding of the OTS-words subregions, suggesting that they are flexibly recruited for processing any kind of readable stimuli.

## 2. Materials and Methods

### 2.1 Participants

18 right-handed volunteers (11 female, 7 male, mean age 28.4 years, SD = 12.8 years) were recruited from Stanford University and surrounding areas and participated in two experimental sessions conducted on different days. Two participants had to be excluded due to excessive head motions (more than 3 voxel within and 3.5 voxels between scan motion), and one participant was excluded due to an incidental finding. The final data set contained 15 subjects (8 female, 7 male). Subjects had normal or corrected to normal vision and gave their informed written consent. The Stanford Internal Review Board on Human Subjects Research approved all procedures.

### 2.2 Data acquisition and preprocessing

#### 2.2.1 Acquisition

Data was collected at the Center for Cognitive and Neurobiological Imaging at Stanford University, using a GE 3 tesla SIGNA Scanner with a 32-channel head coil. 48 slices were acquired, covering the occipitotemporal and most of the frontal cortex using a T2*-sensitive gradient echo sequence (resolution: 2.4 mm × 2.4 mm × 2.4 mm, TR: 1000 ms, TE: 30 ms, FoV: 192 mm, flip angle: 62°, multiplexing factor of 3). A whole-brain, anatomical volume was acquired as well, once for each participant, using a T1-weighted BRAVO pulse sequence (resolution: 1mm × 1 mm × 1 mm, TI=450 ms, flip angle: 12°, 1 NEX, FoV: 240 mm).

#### 2.2.2 Preprocessing

The anatomical brain volume of each participant was segmented into grey and white matter using FreeSurfer (http://surfer.nmr.mgh.harvard.edu/). Manual corrections were made using ITKGray (http://web.stanford.edu/group/vista/cgi-bin/wiki/index.php/ItkGray), and each subject ‘s cortical surface was reconstructed. Functional data was analyzed using the mrVista toolbox (http://github.com/vistalab) for Matlab. The fMRI data from each experiment was motion-corrected, both within and between runs, and then aligned to the anatomical volume. To increase precision, all analyses were conducted in the native brain space of each participant, and no spatial smoothing was applied. The time course of each voxel was high-pass filtered with a 1/20 Hz cutoff and transformed from arbitrary units to percentage signal change. For each experiment, we created a separate design matrix and convolved it with the hemodynamic response function (HRF) implemented in SPM (http://www.fil.ion.ucl.ac.uk/spm) to generate predictors for each experimental condition. Regularized response coefficients (betas) were estimated for each voxel and each predictor using a general linear model (GLM) indicating the magnitude of response for that condition.

#### 2.3 Experimental Design

#### 2.3.1 Experiment 1: Localizer

The first experiment was designed to detect voxels which show higher responses for text compared to other visual stimuli, as in previous work (Stigliani et al., 2015).

##### Stimuli

Stimuli were images belonging to one of five different categories: text, faces, limbs, objects, and houses. In order to also detect color-sensitive voxels, the stimuli were presented either in grayscale or in color.

##### Trial structure

All stimuli were shown for 500 ms, in blocks of 8 stimuli. Each participant completed 4 runs; each run was 5 minutes long and included 10 repetitions of each stimulus category.

##### Task

Participants were asked to fixate a black dot in the center of the screen and to indicate immediate repetitions of a stimulus by pressing a button (1-back task, Fig 1d). For further details on the experimental design see Stigliani et al. (2015).

#### 2.3.2 Experiment 2: fMRI-adaptation

The goal of the second experiment was to leverage fMRI adaptation effects to probe stimulus preferences in the previously identified OTS-words and control regions, with high spatial accuracy. *Stimuli*: Stimuli consisted of two pairs of images, presented at the center of the screen, whereas each pair formed an English compound word (e.g., raincoat). The compound words were presented in three different formats: as emojis, as text, or in a mixed format, in which one pair was an emoji and one pair was text. All stimuli were either white or black on a gray background.

##### Trial structure

In each trial, two image pairs were presented, which were either repeated (e.g., “raincoat” followed by “raincoat”) or non-repeated (e.g., “raincoat” followed by “sunflower”). Each pair was presented for 1s (Fig 1e). Participants completed 6 runs; each run had a duration of ∼5 minutes and contained 72 experiment trials. Trial conditions were balanced in regards to stimulus format and repetition within each run.

##### Task

The image pairs were presented either in black or white in ∼50% of the trials. Subjects had to press a button whenever they saw a white image pair (black/white detection).

#### 2.3.3 Experiment 3: Task and stimulus preferences

The goal of experiment 3 was to probe the influence of different tasks (reading task vs. color judgment task) and stimuli (emoji vs. text stimuli) on neural responses in the OTS-words subregions.

##### Stimuli

Participants were presented with two stimuli shown consecutively, each stimulus depicting an English word. The stimuli were either both in text or emoji format. Each stimulus pair either formed a meaningful English compound (sun + flower) or a meaningless term (flower + sun). All pairs were presented in color on a gray background. The pairs had either the exact same color or similar, yet distinguishable, hues (Fig 1f). Behavioral pilot experiments performed on Amazon Mechanical Turk (MTurk) and in the lab were used to i) ensure that the compound words are easily identifiable even in emoji format and, ii) titrate the hues used in the color task, until task difficulty was well-matched between the color and the reading task.

##### Trial structure

At the beginning of a trial, participants were given a cue indicating which task should be performed (“Read” or “Color”), which was shown for 1s. After the cue, the two images were shown consecutively, each for 1s, followed by an answer screen presented for 2s (Fig 1f, bottom). Subjects completed 8 runs, each lasting ∼5 minutes and containing 24 trials. Stimulus and task conditions were balanced across runs so that each compound word came up equally often in each experimental condition.

##### Task

The experiment included two tasks, a reading, and a color judgment task. When performing the reading task, participants had to read the stimuli and decide whether they form a meaningful English compound word (match) or not (mismatch). The color judgment task required participants to carefully compare the hues of the two presented stimuli, and to decide whether they are the same (match) or slightly different (mismatch). Participants responded by selecting “match” or “mismatch” on the answer screen via a button press, whereas the correct answer was presented randomly either on the left or the right side of the answer screen.

### 2.4 Regions of Interest Definition

Functional Regions of Interest (fROIs) were defined in the native brain space of each participant, using both functional and anatomical criteria. We used all runs from the first experiment (localizer) to define the fROIs. The fROIs defined during experiment 1 were then used to evaluate responses in the other two experiments. We did not apply any spatial smoothing and avoided group-based analyses, as those increase the risk of spurious overlap between closely neighboring fROIs (Weiner & Grill-Spector, 2013). We defined two sets of fROIs in the ventral temporal cortex:

#### 2.4.1 OTS-words

We localized the left OTS-words subregions by contrasting responses to text stimuli with responses to other visual categories (limbs, faces, objects, and houses). Consistent with prior results (Lerma-Usabiaga et al., 2018), we found two self-contained regions in the left middle and posterior occipitotemporal sulcus (OTS) that showed a preference for text over other visual stimuli (T=3, voxel level, uncorrected). We refer to them as middle OTS-words (mOTS-words) and posterior OTS-words (pOTS-words). We were able to define pOTS-words in all 15 participants. On average, the size of pOTS-words was 722.8 mm^3^ (SD = 459.9 mm^3^, N = 15). We were also able to identify mOTS-words in all 15 subjects. The average size of mOTS-words was 444.2 mm^3^ (SD = 350.7 mm^3^, N=15).

#### 2.4.2 Color-sensitive region

In addition to the OTS-words subregions, we defined three color-sensitive patches along the left midfusiform sulcus, containing voxels which showed significantly higher responses for colored compared to grayscale images (T=3, voxel level, uncorrected) (Lafer-Sousa et al., 2016). We refer to them as the anterior, central, and posterior color regions. The anterior color region (Ac-Color) could be identified in all 15 subjects in the left hemisphere, with an average size of 103.5 mm^3^ (SD = 95.5 mm^3^, N = 15). The left posterior color region (Pc-Color) could also be defined for all 15 subjects with an average size of 384 mm^3^ (SD = 228.6 mm^3^, N = 15). A left central color region, Cc-color, could be defined for all 15 subjects, spanning 409.4 mm^3^ (SD = 238.7 mm^3^, N = 15). Since Cc-color most closely neighbors the OTS-words subregions, we chose this region as the control fROI.

Additionally, we also defined constant size 7.5 mm disk fROIs for mOTS-words, pOTS-words, and Cc-color in the left hemisphere, in order to match fROI sizes across participants for the multivoxel pattern analyses (MVPAs, see below). These disk fROIs were centered on the respective region, and the radius of the disk was chosen to match the average size of the OTS-words subregions and Cc-color.

### 2.5 Statistical analysis

#### 2.5.1 Univariate Analysis

We first extracted the average time course in percentage signal change for each condition from mOTS-words, pOTS-words, and the control region. Then we applied a general linear model (GLM) to estimate betas, indicating the magnitude of response for each condition. Consequently, the bar graphs in Figures 2, 4, and 6 show betas in units of % signal change ± SEM. Importantly, we used independent data for fROI definition (experiment 1) and signal extraction (experiments 2 and 3). We conducted repeated measures analyses of variance (rmANOVAs) on the betas from each fROI. **Exp2:** rmANOVAs used stimulus (emoji, text, mixed) and repetition (repeated vs. non-repeated) as factors; **Exp3:** rmANOVAs used stimulus (emoji or text) and task (reading or color judgment) as factors. We also conducted rmANOVAs for both **Exp 2** and **Exp 3** where we added ROI (mOTS-words, pOTS-words) as an additional factor to compare responses across the two OTS-words subregions. Post-hoc tests were applied, when the rmANOVAs revealed significant main effects or interaction effects between factors. Where applicable, we used Bonferroni correction for multiple comparisons.

**Figure 2.**
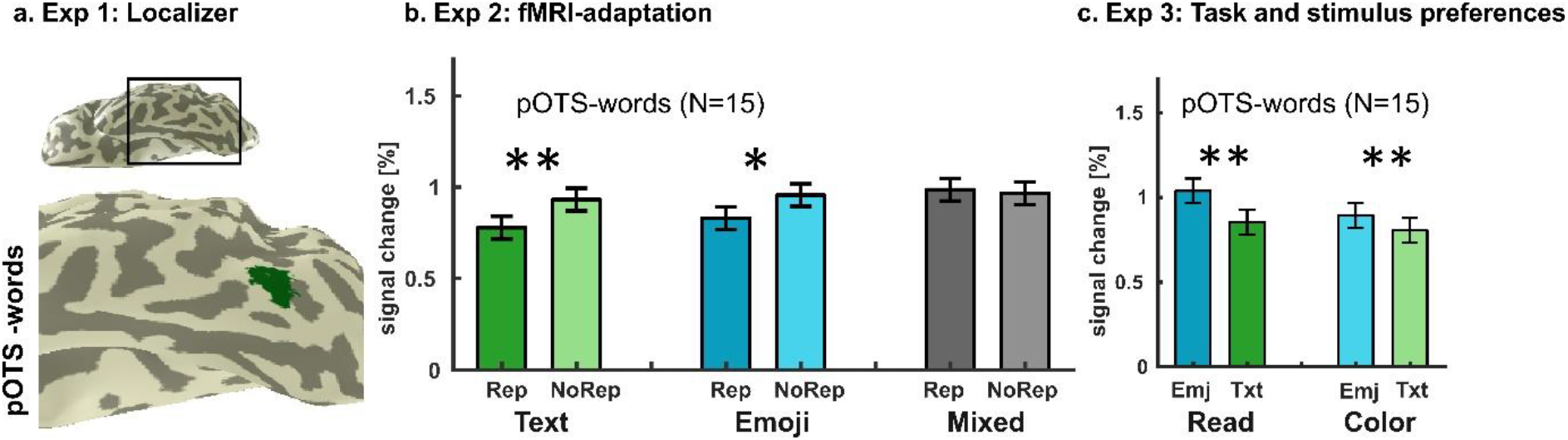
pOTS-words shows fMRI adaptation for both emoji and text stimuli in Exp 2 and an overall preference for emojis in Exp 3. **a. Exp1: Localizer:** Location of pOTS-words in the left hemisphere of a representative subject. pOTS-words was identified as a cluster of voxels in the posterior occipitotemporal sulcus that responded higher to text than limbs, objects, houses, and faces (T = 3, voxel level, uncorrected). **b. Exp 2: fMRI-adaptation**: Mean responses ±SEM of pOTS-words across all subjects (N=15). A main effect of repetition and an interaction between repetition and stimulus was observed. Post-hoc tests revealed significant fMRI adaptation for text and emoji stimuli, but not the mixed condition. Stars indicate post-hoc results, showing significant differences between repeated and non-repeated trials; * p <0.05, ** p < 0.01. **c. Exp 3: Task and stimulus preferences**. Mean responses *±* SEM of pOTS-words across all subjects (N=15). A main effect of stimulus was observed, whereas post-hoc tests showed higher responses for emoji than text stimuli for both tasks. Stars indicate post-hoc results; ^*^ p <0.05, ^**^ p < 0.01, ^***^ p<0.001. *Abbreviations*: Rep = repeated trials, NoRep = non-repeated trials, Emj = emoji trials, Txt = text trials.

#### 2.5.2 Multivoxel Pattern Analysis (MVPA)

We conducted MVPAs (Haxby et al., 2001)on distributed responses across the 7.5 mm disk fROIs for mOTS-words, pOTS-words, and Cc-color. A GLM was calculated to estimate the responses to each experimental condition separately for each voxel. The responses were then normalized by subtracting each voxel ‘s mean response and z-standardized.

For each experiment and each condition, we calculated the correlation among each pair of multivoxel patterns (MVPs), using a leave-one-run-out procedure. These correlations were summarized in representational similarity matrices (RSMs). By using the leave-one-run-out procedure as cross-validation, we were able to evaluate the reliability of the observed MVP across runs. The RSMs were calculated for each subject separately and then averaged across subjects. In addition to that, we trained winner-takes-all (WTA) classifiers separately on the responses of experiments 2 and 3, to observe which experimental conditions can be decoded from the MVPs of each fROIs. In experiment 2, classifiers were initially trained to decode stimulus (emoji, text, mixed) and repetition (repeated vs. non-repeated). However, as the mixed condition includes both emoji and text stimuli, the classifier could not learn patterns associated with each stimulus format when this condition was included. We therefore reran the MVPAs for experiment 2 excluding all mixed trials. For experiment 3, WTA-classifiers were trained to decode stimulus (emoji vs. text) and task (reading task vs. color judgment). All classifiers used a leave-one-run-out procedure, where the training set consisted of all runs but one, and the test set included the left-out run. Results report the aggregated classifier performances across all iterations of leave-one-run-out combinations. To quantify the classifiers ‘ performance, we computed paired t-tests against chance level for stimulus and task decoding. Where applicable, Bonferroni correction for multiple comparisons was applied.

We also created RSMs combining responses from experiments 2 and 3, to test MVPs for stimulus (emoji vs. text) and tasks (reading, color judgment, black/white judgment) across experiments. We included all conditions from experiment 3, and all emoji and text trials from experiment 2, averaging repeated and non-repeated trials. We normalized the data by subtracting the mean response separately for each experiment before merging them, as the average response levels differed between experiments. As experiments 2 and 3 also differed in their number of runs, a leave-one-run-out procedure was not feasible, which is why we divided the data set into even and odd runs for the RSA. RSMs were calculated for each subject separately and then averaged across subjects.

Finally, we conducted additional rmANOVAs on the WTA classifier decoding accuracy with decoding type (**Exp 2**: stimulus and repetition, **Exp 3**: stimulus and task) and ROI (mOTS-words, pOTS-words) as factors, to compare decoding performance across the OTS-words subregions. Post-hoc tests were applied, when the rmANOVAs revealed significant main effects or interaction effects between factors. Where applicable, we used Bonferroni correction for multiple comparisons.

#### 2.5.4 Analysis of behavioral responses during fMRI

We calculated each subject ‘s average performance (measured in % correct *± SE*) and reaction times separately for each experiment. Further, we tested for significant performance differences between experimental conditions, by conducting rmANOVAs on the behavioral responses. **Exp1:** rmANOVAs used stimulus (words, limbs, objects, houses, faces) and color (colored vs. gray-scale) as factors; **Exp2:** rmANOVAs used stimulus (emoji, text, mixed) and repetition of stimulus (repeated vs. non-repeated) as factors; **Exp3**. rmANOVAs used stimulus (emoji or text) and task (reading or color judgment) as factors. Post-hoc tests were applied, when the rmANOVAs revealed significant main effects or interaction effects between factors. Where applicable, Bonferroni correction for multiple comparisons was applied.

### 2.6 Code and Data availability

The fMRI and T1w-anatomical data were analyzed using the open-source mrVista software (available in GitHub: http://github.com/vistalab/vistasoft) and FreeSurfer (available at https://surfer.nmr.mgh.harvard.edu/) respectively. Source data and code for reproducing all figures and statistics are made available in GitHub as well (https://github.com/EduNeuroLab/read_emojis_ots). The raw data collected in this study will be made available by the corresponding author upon reasonable request.

## 3. Results

### 3.1 Task difficulty was well-matched across conditions in each of the three experiments

The goal of the first experiment was to localize mOTS-words and pOTS-words in each participant by contrasting responses to text stimuli with those elicited by other visual stimulus categories. Subjects (N=15) were presented with images of texts, limbs, faces, objects, and houses while they performed a 1-back task. We used rmANOVAs to compare performance and reaction time across conditions. On average, participants ‘ accuracy (± SE) was 60% (*±*4 %). There was no significant difference in accuracy between the different stimulus categories (F(4,11) = 0.45, p = 0.77) or between colored or grayscale images (F(1,14) = 0.81, p = 0.38) and no interaction between stimulus and color (F(4,11) = 1.68, p = 0.17). Participants ‘ reaction time (*±* SE) was on average 260 ms (*±* 40 ms) and did not differ between stimulus categories (F(4,11) = 0.47, p = 0.76) or between colored or grayscale images (F(1,14) = 2.13, p = 0.12). There was also no interaction between stimulus and color (F(4,11) = 0.53, p = 0.71).

The goal of the second experiment (fMRI-adaptation) was to test whether there is a difference in neural adaptation for emojis compared to text stimuli in mOTS-words and pOTS-words. The images were shown either in white or black and the participant ‘s task was to press a button when white images were shown. We tested for differences in subjects ‘ performance (*±* SE) as well as in their reaction times (*±* SE) by conducting rmANOVAs of behavioral responses with repetition and stimulus as factors. On average, participants responded correctly in 91% (*±* 1%) of the trials. Participants ‘ performance did not significantly differ between emoji, text, and mixed trials (F(2,13) = 0.40, p = 0.68) or between repeated and non-repeated trials (F(1,14) = 0.20, p = 0.66). We found no interaction effect between stimulus and repetition (F(2,13) = 0.70, p = 0.55). The subjects ‘ reaction time (± SE) was on average 304 ms (± 10ms). We found no significant differences in response time between emoji, text, and mixed trials (F(2,13) = 0.99, p = 0.38) or between repeated and non-repeated trials (F(1,14) = 0.10, p = 0.75) and no interaction between stimulus and repetition (F(2,13) = 0.69, p = 0.5).

The third experiment aimed to uncover the influence of different tasks (reading vs. non-reading) and stimuli (text vs. emoji) on responses in mOTS-words and pOTS-words. We performed rmANOVAs, using task and stimulus as factors to evaluate performance and reaction times. On average, subjects ‘ performance (± SE) was 95 % (± 2%) correct. There were no effects of task (F(1,14) = 0.33, p = 0.67) or stimulus (F(1,14) = 0.22, p = 0.65) and no interaction effect between stimulus and task (F(1,14) = 0.21, p = 0.65) on participants ‘ accuracy. Participant ‘s average reaction time (± SE) was 669 ms (± 25ms). We found a significant main effect of stimulus (F(1,14) = 8.17, p = 0.01) and task (F(1,14) = 6.50, p = 0.02) and a significant interaction between stimulus and task (F(1,14) = 7.99, p = 0.01) on participant ‘s reaction times. Post-hoc analyses revealed significantly longer reaction times for emoji compared to text stimuli in the reading task (p = 0.0004).

### 3.2 pOTS-words shows higher responses for emoji stimuli than text stimuli independent of the task

We identified pOTS-words in the left hemisphere in all 15 participants by contrasting responses to text stimuli with responses elicited by other visual categories (faces, objects, limbs, and houses) (Fig 2a shows an example pOTS-words in a representative subject). To test if pOTS-words is only responsive towards text stimuli or also other, readable stimuli, such as emojis, in experiment 2 we leveraged an fMRI-adaptation design. We found no significant main effect of stimulus (F(2,13) = 1.53, p = 0.23), but a main effect of repetition (F(1,14) = 14.99, p = 0.002), as well as an interaction between repetition and stimulus (F(2,13) = 5.30, p = 0.01). Post-hoc tests revealed a significant effect of repetition for both text (p = 0.001) and emoji stimuli (p = 0.03), but not for the mixed condition (p= 0.39) (Fig 2b).

In experiment 3, we further probed task and stimulus preferences in pOTS-words by manipulating the participant ‘s tasks and the presented stimuli orthogonally. pOTS-words showed a main effect of stimulus (F(1,14) = 12.37, p = 0.003), but no significant main effect of task (F(1,14) = 3.85, p = 0.07) and no significant interaction effects (F(1,14) = 1.91, p = 0.18) (Fig 2c). Post-hoc tests revealed higher responses for emoji stimuli than text stimuli in pOTS-words (p = 0.003).

### 3.3 Distributed responses across pOTS-words encode the visual stimulus

In addition to the univariate results presented above, we also probed what information can be decoded from distributed responses within pOTS-words. For this, we performed MVPA within 7.5mm disk ROIs centered on pOTS-words. We used a constant disk ROI to equate the number of voxels used in the MVPA across participants.

In experiment 2, we generated an RSM comparing repeated and non-repeated emoji and text trials using a leave-one-run-out-approach. Thus, the diagonal of the RSM indicates similarity with the same condition across runs, and the off diagonal indicates similarity among different conditions across runs. The RSM reveals positive correlations for the same condition across runs as well as for the same stimuli across repeated and non-repeated trials (Fig 3a). To quantify if information about stimulus and/or repetition can be successfully decoded from the distributed responses in pOTS-words, we trained WTA (winner-takes-all) classifiers. We found that the stimulus but not repetition can be successfully decoded from distributed pOTS-word responses. Classifier performance reached an accuracy (*±* SEM) of 86.28% (*±*4.72%) for text and 88.33% (*±*3.93%) for emoji stimuli, which in both cases significantly exceeded 50% chance level (text: p< 0.0001, emojis: p < 0.0001, significant with Bonferroni corrected threshold of p<0.01). Classifier performances for distinguishing between repeated and non-repeated trials reached an accuracy (± SEM) of 44.22% (±4.76%) for repeated and 57.11% (±4.30%) for non-repeated trials, which did not exceed chance level (repeated trials: p = 0.88, non-repeated trials: p = 0.06) (Fig 3b). Accordingly, a direct comparison showed that the classifier decoding stimulus yielded a significantly higher performance than the classifier differentiating repeated and non-repeated trials (p < 0.0001).

**Figure 3.**
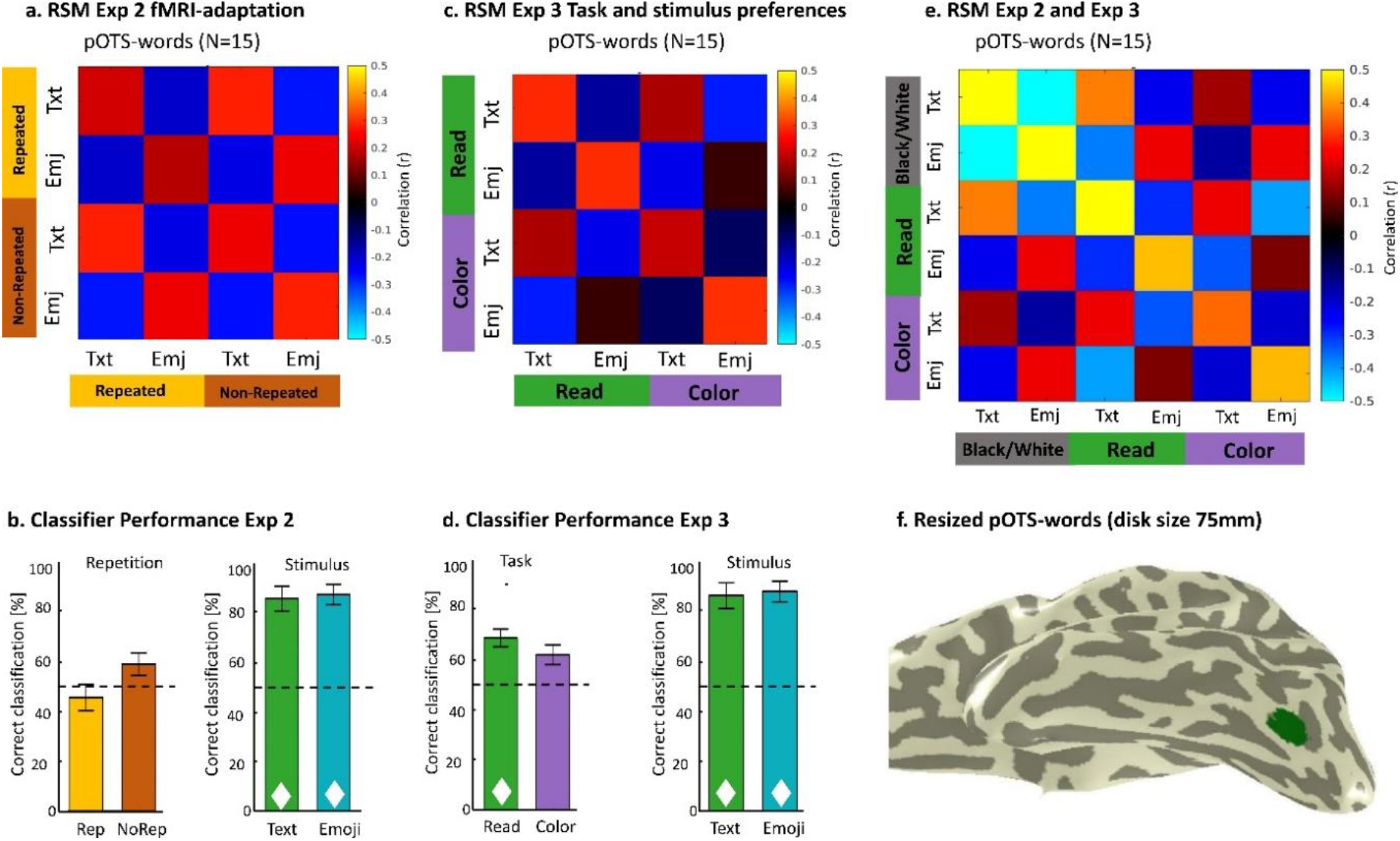
Distributed responses of pOTS-words encode different stimuli, both within and across experiments. **a**. RSM for experiment 2 data from pOTS-words in the left hemisphere across all subjects. Conditions are arranged by stimulus (text vs. emoji) and grouped by repetition (repeated vs. non-repeated). **b**. Mean ± SEM WTA classification performance in experiment 2 for stimulus and repetition. Classifier performance significantly exceeded 50% chance level for decoding both stimulus categories but not repetition. **c**. RSM for experiment 3 data from pOTS-words in the left hemisphere across all subjects. Conditions are arranged by stimulus (text vs. emoji) and grouped by task (reading vs. color). **d**. Mean ± SEM of WTA classification performance in experiment 3 for stimulus and task. Classifier performance for decoding the reading task and for decoding both stimulus categories significantly exceeded chance level. **e**. Joined RSM for both experiments 2 and 3 data from pOTS-words in the left hemisphere across all subjects. Conditions are arranged by stimulus (text vs. emoji) and grouped by task (reading, color judgment, black/white detection). **f**. Disk fROI centered on pOTS-words is shown in a representative subject and was used for all MVPA analyses. *Dotted line*: chance level (50%); ✦ classifier performance significantly above chance level with a Bonferroni corrected threshold of p <0.01; *Abbreviations*: RSM=representational similarity matrix, WTA=winner-takes-all, Rep: repeated trials, NoRep = non-repeated trials, Emj = emoji trials, Txt = text trials.

In experiment 3, we created an RSM comparing stimuli (text, emoji) and tasks (reading, color, Fig 3c) again using a leave-1-run-out approach. As Fig 3c shows, the RSM revealed positive correlations for the same stimulus under different tasks. In addition, WTA classifiers were trained separately for stimulus and task, to determine which kind of information (stimulus, task, or both) can be successfully decoded from distributed pOTS-words responses. Classification performances (*±* SEM) for decoding text stimuli (85.42% *±* 4.88%) and emoji stimuli (87.92% *±* 4.30%) were significantly higher than 50% chance level (emoji trials: p<0.0001, text trials: p<0.0001, significant with Bonferroni corrected threshold of p<0.01). When differentiating between tasks, the classifier ‘s performance (± SEM) was on average at 65%. Its performance (*±* SEM) was 68.54 % (*±*3.50%) for reading and 61.88% (±3.92%) for color judgments, with decoding accuracy for reading significantly exceeding 50% chance level (reading: p = 0.008, significant with Bonferroni corrected threshold of p<0.01; color judgment: p = 0.02, not significant with Bonferroni corrected threshold of p<0.01) (Fig 3d). A direct comparison showed that decoding accuracy for decoding the stimulus was significantly higher than for decoding the task (p < 0.0001). These results suggest that while both the stimulus and the reading task are decodable from distributed responses in pOTS-words, pOTS-words contains more information regarding the visual stimulus than the task.

Finally, we merged data from experiments 2 and 3 to further probe distributed responses elicited by different stimuli and tasks across experiments. For experiment 2 we averaged across repeated and non-repeated trials. The resulting RSM shows that MVPs for the same stimulus (emoji and text) are positively correlated across experiments, suggesting that these stimuli induce stable and reproducible MVPs across the three tasks (reading, color detection, black/white detection). In contrast, there is no similarity in responses across stimuli within a task. Thus, our results suggest that pOTS-words is predominantly involved in encoding stimuli not tasks.

### 3.4 mOTS-words shows higher responses for emoji than text stimuli during the reading task

From experiment 1 data, we identified mOTS-words in each participant ‘s native brain space by contrasting responses to text stimuli with those evoked by other visual stimuli. mOTS-words was successfully identified in all 15 subjects in the left hemisphere. As an example, mOTS-words from a representative subject is shown in Fig 4a. To test if mOTS-words is sensitive only to text or also to other readable visual stimuli, such as emojis, we examined the data from the fMRI adaptation experiment and conducted rmANOVAs with factors of repetition (repeated, non-repeated) and stimulus format (emoji, text, mixed). mOTS-words did not show significant effects of repetition (F(1,14) = 1.12, p = 0.31) or stimulus format (F(2,13) = 0.90, p = 0.42) (Fig 4b). Further, using the data from the third experiment, we tested task and stimulus preferences in mOTS-words. rmANOVAs from mOTS-words with task (reading, color) and stimulus (text, emoji) as factors showed a main effect of stimulus (F(1,14) = 7.54 p = 0.02) and a main effect of task (F(1,14) = 13.28, p = 0.003) (Fig 4c) as well as an interaction between task and stimulus (F(1,14) = 7.66, p = 0.02). Post-hoc t-tests revealed higher responses for emoji than text stimuli during the reading task (p = 0.0005) as well as higher responses for the reading compared to the color task (p=0.003).

**Figure 4.**
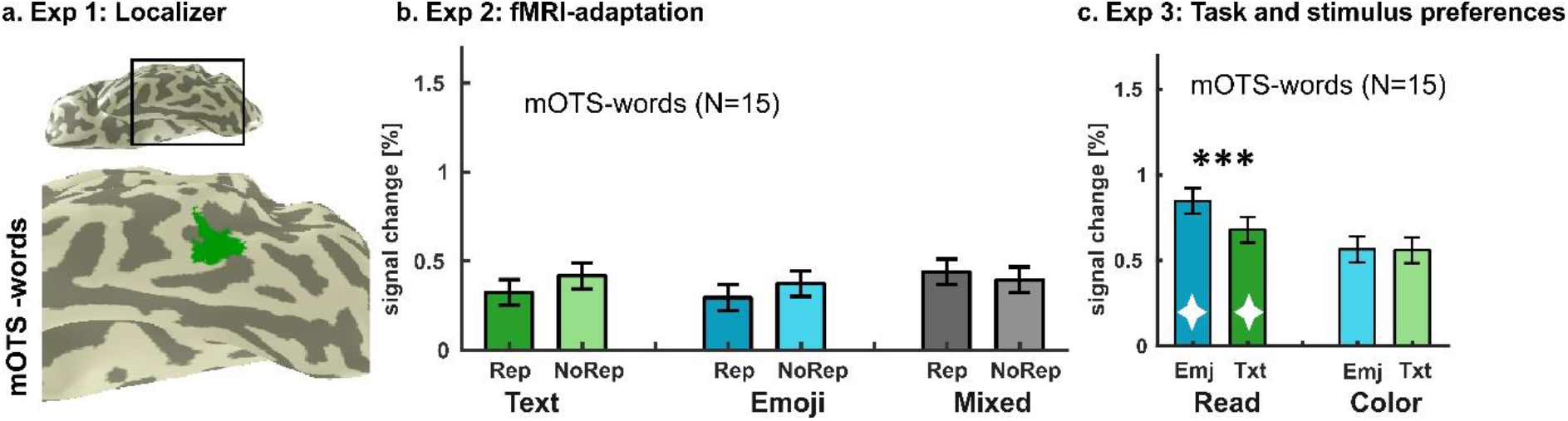
mOTS-words shows higher responses for emoji than text stimuli during a reading task. **a. Exp 1: Localizer**. Location of mOTS-words in the left hemisphere of a representative subject. mOTS-words was identified as a cluster of voxels that respond higher to text than limbs, objects, houses, and faces (T = 3, voxel level, uncorrected). **b. Exp 2: fMRI-adaptation**: Mean responses of mOTS-words across all subjects (N=15) ±SEM. No significant effects of repetition or stimulus were detected. **c. Exp 3: Task and stimulus preferences**. Mean responses ±SEM of mOTS-words across all subjects (N=15). We found a main effect for task and stimulus as well as an interaction effect between task and stimulus. Post-hoc tests revealed a preference for emoji stimuli during the reading task. Stars indicate results of post-hoc tests, showing significantly higher responses for emoji stimuli during a reading task, ^*^ p <0.05, ^**^ p < 0.01,^***^ p<0.001; diamonds indicate significantly higher responses for one task vs. the other task, p<0.05. *Abbreviations*: Rep: repeated trials, NoRep = non-repeated trials, Emj = emoji trials, Txt = text trials.

### 3.5 Distributed responses across mOTS-words encode stimulus and task

In addition to the univariate results presented above, we also probed what information can be decoded from distributed responses within 7.5mm disk ROIs centered on mOTS-words.

In experiment 2, we calculated a RSM comparing MVPs of repeated and non-repeated emoji and text trials. We observed high MVP correlations for the same stimuli across repeated and non-repeated trials (Fig 5a). Further, WTA classifiers performances (*±* SEM) reached 79.94% (*±*3.22%) accuracy for text and 77.83% (±4.55%) accuracy for emoji stimuli, which in both cases significantly exceeded 50% chance level (text : p<0.0001, emojis: p<0.0001, significant with Bonferroni corrected threshold of p<0.01) (Fig 5b). Classifier performances (±SEM) for distinguishing between repeated and non-repeated trials was at 48.94% (±3.40%), for repeated and at 56.06% ±(3.11%) for non-repeated trials, which did not exceed 50% chance level (repeated: p=0.62, non-repeated: p=0.04, not significant with Bonferroni corrected threshold of p<0.01). Accordingly, the classifier decoding stimulus performed significantly better than the one decoding repetition (p<0.0001).

**Figure 5.**
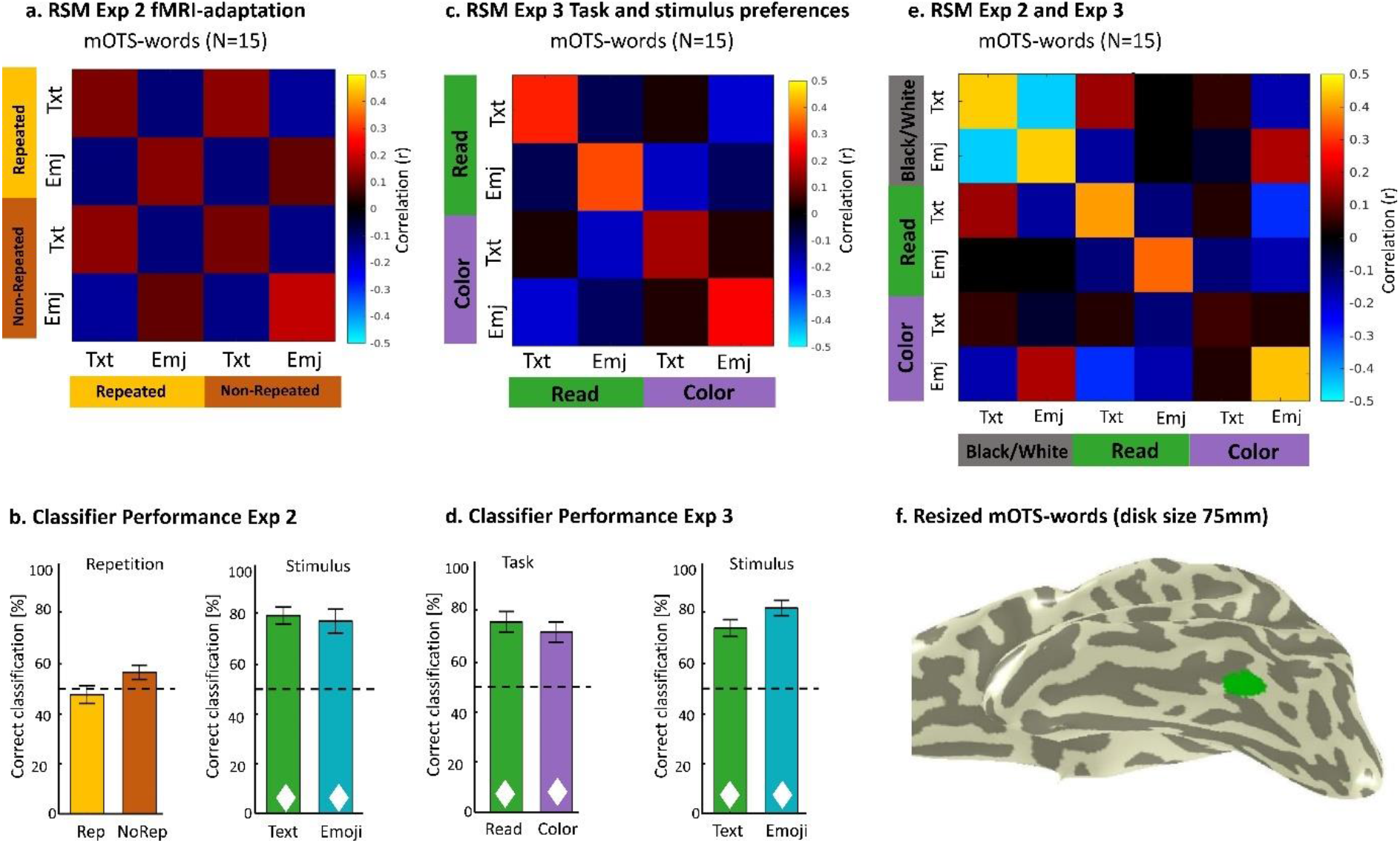
Distributed responses of mOTS-words encode stimuli and tasks, both within and across experiments. **a**. Mean RSM for experiment 2 for mOTS-words in the left hemisphere across all subjects. Conditions are arranged by stimulus (text vs. emoji) and grouped by repetition (repeated vs. non-repeated trials). **b**. Mean ± SEM WTA classification performance in experiment 2 for stimulus and repetition. Classifier performance significantly exceeded 50% chance level for decoding both stimuli but not decoding repetition. **c**. Mean RSM for experiment 3 for mOTS-words in the left hemisphere across all subjects. Conditions are arranged by stimulus (text vs. emoji) and grouped by task (reading vs. color judgment). **d**. Mean ± SEM WTA classification performance in experiment 3 for stimulus and task. Classifier performance was significantly above 50% chance level for decoding both stimuli and tasks. **e**. Mean RSM across experiments 2 and 3 for mOTS-words in the left hemisphere across all subjects. Conditions are arranged by stimulus (text vs. emoji) and grouped by task (black/white detection vs. reading vs. color judgment). **f**. Disk fROI centered on mOTS-words is shown in a representative subject and was used for all MVPA analyses. *Dotted line*: chance classification level (50 %); ✦ classifier performance significantly above chance level with a Bonferroni corrected threshold of p<0.01; *Abbreviations*: RSM=representational similarity matrix, WTA=winner-takes-all, Rep: repeated trials, NoRep = non-repeated trials, Emj = emoji trials, Txt = text trials.

In experiment 3, we created a RSM comparing MVPs across stimuli (emoji, text) and tasks (reading, color judgment) (Fig 5c). The RSM shows high MVP correlations only along the diagonal, suggesting that MVPs in mOTS-words distinguish between both stimuli and tasks. To further quantify these effects, we trained WTA classifiers, separately for stimulus and task, to determine which kind of information (stimulus, task, or both) can be successfully decoded from MVPs in mOTS-words (Fig 5d). The classifier ‘s accuracy (*±* SEM) in % for decoding the reading task was 78.54% (*±*4.17%) and for decoding the color task was 74.38% (±4.06%), which in both cases, was significantly higher than chance level (reading task: p<0.0001, color task: p<0.0001, significant with Bonferroni corrected threshold of p<0.01). The classifier trained to differentiate between emoji and text stimuli had an accuracy of 80.42% (±4.07%) for decoding emojis and 72.50% (±3.29%) for decoding text stimuli. Classifier performances significantly exceeded chance level for both stimulus conditions (emoji stimuli: p<0.0001, text stimuli: p<0.0001, significant with Bonferroni corrected threshold of p<0.01). We found no significant differences, when we compared the classifier ‘s accuracy for stimulus vs. task decoding (p = 1).

Next, we calculated an RSM combining data from experiments 2 and 3 and compared MVPs for emoji and text trials across three tasks (black/white detection, reading, and color judgment) (Fig 5e). We used all trials from experiment 3 and the emoji and text trials from experiment 2, averaged across repeated and non-repeated trials. The resulting RSM showed high correlations mostly along the diagonal suggesting that changing either task or stimulus generates a distinct pattern of response in mOTS-words.

### 3.6 Cc-color shows higher responses for the color task than the reading task

In order to identify the color patches along the midfusiform sulcus, in experiment 1, we contrasted responses to colored images with gray-scale images. Three color regions were defined. Here we focus on Cc-color, as it was successfully identified in all 15 subjects and is adjacent to both OTS-words subregions in the left hemisphere. Cc-color was used as a control region to evaluate the specificity of the response patterns observed in mOTS-words and pOTS-words (Fig 6a).

**Figure 6.**
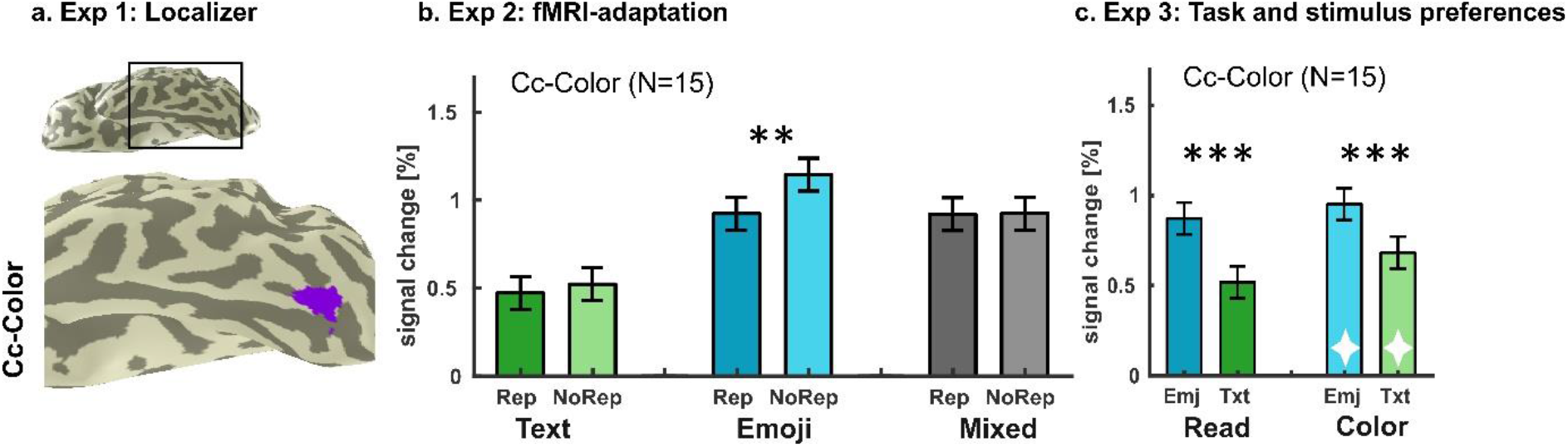
The central color region shows preferences for emojis and the color judgment task. **a. Exp 1: Localizer**: Location of Cc-color in the left hemisphere of a representative subject. Cc-color was identified as voxels medial to the mfusiform gyrus that responded higher to colored compared to grey-scale images (T = 3, voxel level, uncorrected). **b. Exp 2: fMRI-adaptation:** Mean responses ±SEM of Cc-color across all subjects (N=15). Main effects of stimulus and repetition as well as an interaction between stimulus and repetition were observed. Post-hoc tests showed significant fMRI adaptation only for emoji stimuli. Stars indicate results of post-hoc tests, showing significant differences between repeated and non-repeated trials for the emoji condition; * p <0.05, ^**^ p < 0.01. **c. Exp 3: Task and stimulus preferences**. Mean responses ±SEM of Cc-color across all subjects (N=15) in experiment 3. A main effect of stimulus was observed, with higher responses for emoji than text stimuli for both tasks. Stars indicate post-hoc results, ^*^ p <0.05, ^**^ p < 0.01,^***^ p<0.001; diamonds indicate significantly higher responses for one task vs. the other task, p <0.05. *Abbreviations*: Exp: experiment, Rep: repeated trials, NoRep = non-repeated trials, Emj = emoji trials, Txt = text trials

In experiment 2, we used fMRI adaptation to probe stimulus representations in Cc-color. In the left hemisphere, we found a main effect of stimulus (F(2,13) = 32.98, p<0.0001) and a main effect of repetition (F(1,14) = 5.37, p = 0.04). Furthermore, we found a significant interaction between repetition and stimulus (F(2,13) = 4.11, p = 0.03) (Fig 6b). Post-hoc tests revealed a significant difference between repeated and non-repeated trials for the emoji stimuli (p = 0.007), but not for the text (p = 0.39) or the mixed condition (p = 0.94).

In experiment 3, where tasks and stimuli were manipulated orthogonally, Cc-color showed a main effect of stimulus (F(1,14) = 81.62, p<0.0001) with a preference for emoji stimuli over text stimuli. Moreover, we found a significant main effect of task (F(1,14) = 20.85, p<0.0001), indicating a preference for the color judgment task over the reading task in Cc-color. This established a double-dissociation of task preferences with mOTS-words (Fig 6c). We found no interaction between task and stimulus (F(1,14) = 2.16, p = 0.16) in Cc-color.

### 3.7 Distributed responses across the central color region encode the visual stimulus

In addition to the univariate results presented above, we also probed what information can be decoded from distributed responses within left Cc-color. For this, we performed MVPA analyses within 7.5mm disk fROIs centered on Cc-color.

In experiment 2, we generated an RSM comparing MVPs across repeated and non-repeated text and emoji trials. The RSM revealed a clear stimulus effect, showing positive correlations between repeated and non-repeated trials when the same stimulus was presented (Fig 7a). We then trained WTA classifiers to test what kind of information (stimulus, repetition, or both) can be successfully decoded from distributed responses in Cc-color (Fig 7b). Classification performance (mean *±* SEM) in % for decoding text yielded 82.67% (*±*5.61%) accuracy and accuracy for decoding emojis was 80.72% (±5.62%). In both cases, decoding accuracy was significantly above 50% chance level (text: p<0.0001, emojis: p<0.0001, significant with Bonferroni corrected threshold of p<0.01). Classification performance (±SEM) in % for detecting repeated trials was 52.50% (±3.06%) and decoding accuracy was 64.00% (±4.07%) for non-repeated trials. Only when decoding non-repeated trials, classifier performance significantly exceeded 50% chance level with a Bonferroni corrected threshold of p<0.01 (repeated: p = 0.21, non-repeated: p = 0.002). When comparing the average performances for decoding stimulus vs. repetition, we found that the classifier decoding stimulus performed significantly better (p = 0.01).

**Figure 7.**
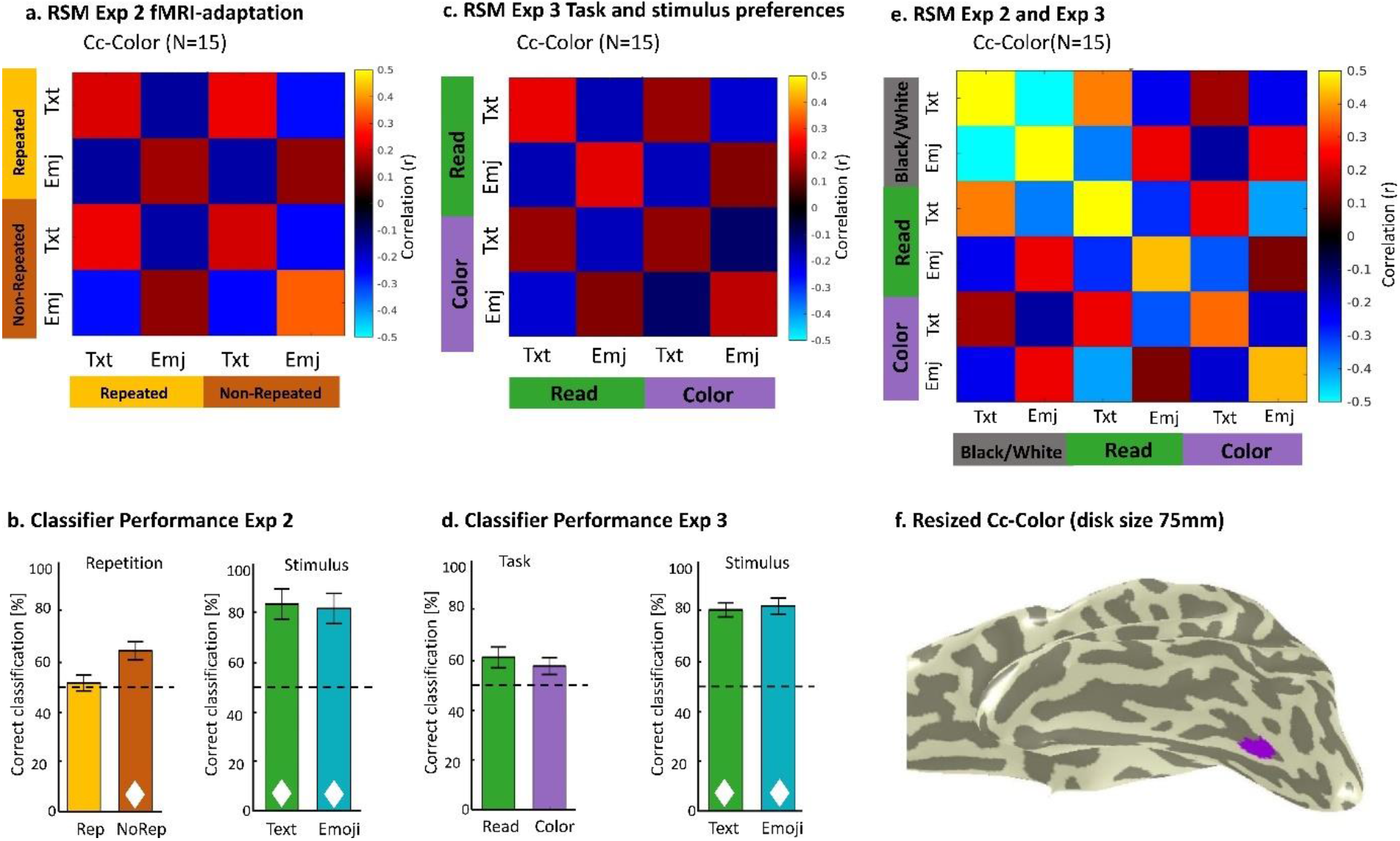
Distributed responses of Cc-color encode visual stimuli, both within and across experiments. **a**. Mean RSM from experiment 2 for Cc-color in the left hemisphere across all subjects. Conditions are arranged by stimulus (text vs. emoji) and grouped by repetition (repeated vs. non-repeated). **b**. Mean±SEM WTA classification performance in experiment 2 for stimulus and repetition. Classifier performance significantly exceeded chance level for decoding both stimulus categories and non-repeated trials. **c**. Mean RSM from experiment 3 for Cc-color in the left hemisphere across all subjects. Conditions are arranged by stimulus (text vs. emoji) and grouped by task (reading vs. color judgment). **d**. Mean±SEM WTA classification performance in experiment 3 for stimulus and task. Classifier performances for decoding stimulus but not task significantly exceeded 50% chance level. **e**. Mean RSM for joined analysis of experiments 2 and 3 for Cc-color in the left hemisphere across all subjects. Conditions are arranged by stimulus (text vs. emoji) and grouped by task (reading vs. color judgment vs. black/white detection). **f**. Disk fROI centered on Cc-color is shown in a representative subject and was used for all MVPA analyses. ✦ classifier performance significantly above chance level with a Bonferroni corrected threshold of p <0.01; *Abbreviations*: RSM: representational similarity matrix, WTA = winner-takes-all, Rep = repeated trials, NoRep = non-repeated trials, Emj = emoji trials, Txt = text trials.

In experiment 3, we created an RSM comparing MVPs across different tasks and stimuli (Fig 7c). Examination of this RSM revealed a strong stimulus effect, as MVPs for the same stimulus under different tasks were positively correlated. Accordingly, WTA classification performances (mean*±*SEM) for emoji trials (81.88±3.30%) and text trials (80.42±2.79%) were significantly higher than 50% chance level (emoji trials: p<0.0001, text trials: p<0.0001, significant with Bonferroni corrected threshold of p<0.01). The classifier ‘s performance (*±*SEM) for distinguishing between the reading and color judgment tasks were 61.46% (*±* 4.18%) for reading and 57.92% (*±*3.39%) for color judgment, neither of which significantly exceeded 50% chance level (reading: p = 0.05, color judgment: p = 0.09, not significant with Bonferroni corrected threshold of p<0.01) (Fig 7d). The average classification performance for decoding the stimuli was significantly higher than for decoding the tasks (p < 0.0001).

Finally, to investigate if similar stimuli induce reproducible MVPs in Cc-color across experiments, we calculated an RSM combining data from experiments 2 and 3. This RSM showed positive correlations for similar stimuli, even across experiments that included different tasks. In sum, these results suggest that MVPs in Cc-color contains information regarding the visual stimulus, but not regarding the task the participants are performing.

### 3.8 Task and stimulus decodability differ between the OTS-words subregions

After examining mOTS-words and pOTS-words separately, we wanted to explicitly test if there are significant differences in task and stimulus selectivity across these regions. To this end, for the univariate data, we conducted additional rmANOVAs adding ROI (mOTS-words, pOTS-words) as a factor. In experiment 2, we found no main effect of stimulus (F(2,13) = 1.56, p = 0.23), but a main effect of repetition (F(1,14) = 5.10, p = 0.04) and ROI (F(1,14) =34.00, p < 0.0001). We also observed a significant interaction between stimulus and repetition (F(2,13) = 4.94, p = 0.01); post hoc tests revealed significant differences between repeated and non-repeated trials for text stimuli only (text: p = 0.008, emoji: 0.09, mixed: 0.23). We found no interaction between repetition and ROI (F(1,14) = 1.88, p = 0.19) and stimulus and ROI (F(2,13) = 0.63, p = 0.58). The three-way interaction of ROI, stimulus and repetition was not significant as well (F(2,13) = 0.17, p = 0.85). In experiment 3, we found a main effect of task (F(1,14) = 11.56, p = 0.004), a main effect of stimulus (F(1,14) = 27.48, p = 0.0001) and a main effect of ROI (F(1,14) = 6.86, p = 0.02). No significant interaction effect between ROI and stimulus (F(1,14)=0.776, p=0.39) was found. The interaction effect between task and stimulus was marginally significant (F(1,14) = 4.53, p = 0.05). The interaction between task and ROI was not significant, although a trend was observed (F(1,14) = 3.35, p = 0.09). The three-way interaction between task, stimulus, and ROI was not significant (F(1,14) = 2.13, p = 0.17).

To further investigate if there are significant differences in the information contained in the distributed responses across the OTS-words subregions, we conducted rmANOVAs on the WTA classifier decoding accuracy with decoding type (Exp 2: stimulus and repetition, Exp 3: stimulus and task) and ROI (mOTS-words, pOTS-words) as factors. In experiment 2, we found a main effect of decoding type (F(1,14) = 47.96, p < 0.0001), but no main effect for ROI (F(1,14) = 1.29, p = 0.28), and no interaction effect (F(1,14) = 3.18, p = 0.10). For experiment 3, we found a main effect of decoding type (F(1,14) = 9.10, p = 0.009), no main effect of ROI (F(1,14) = 0.039, p = 0.8), but a significant interaction between ROI and decoding type (F(1,14) = 11.55, p = 0.004). Post-hoc tests revealed a difference in classifier performance between mOTS-words and pOTS-words for decoding both stimulus (p =0.02) and task (p = 0.02), whereas mOTS-words showed better task decoding and pOTS-words showed better stimulus decoding.

## Discussion

Here we assessed task and stimulus preferences in the OTS-words subregions. In experiment 1, we identified mOTS-words and pOTS-words by contrasting text with other visual stimuli. Experiment 2 showed fMRI-adaptation for text and emoji stimuli in pOTS-words but not in mOTS-words. Finally, in experiment 3, pOTS-words showed a preference for emojis independent of task and mOTS-words showed a preference for emojis during a reading task as well as an overall preference for reading. Corresponding multivariate analyses showed that pOTS-words encodes the visual stimulus, while mOTS-words is sensitive to both stimulus and task.

Our finding that mOTS-words is sensitive to reading while pOTS-words is sensitive only to the visual stimulus aligns with previous studies proposing an increasing processing level along the OTS (Carreiras et al., 2014; Taylor et al., 2019; White et al., 2019; Woolnough et al., 2021). First, while the posterior OTS is proposed to process letters, the anterior OTS is suggested to be responsible for perceiving words as language units (Thesen et al., 2012; Vinckier et al., 2007). Accordingly, the local combination detector model (Dehaene et al., 2005) posits that words are encoded through a posterior-to-anterior hierarchy of neuronal populations, which are tuned to process increasingly more complex features of words. Indeed, recent work suggests that the more posterior pOTS-words is involved in visual processing, while mOTS-words plays a role in integrating vision and language (Lerma-Usabiaga et al., 2018), which is supported by differential structural (Lerma-Usabiaga et al., 2018) and functional connectivity (Yablonski et al., 2023) of the sub-regions. This increasing processing level also appears to be functionally relevant, as impairments in the OTS-words processing hierarchy have been associated with dyslexia (van der Mark et al., 2009). Together, these differences in processing level suggest that mOTS-words and pOTS-words should be evaluated separately. For this, it is helpful to perform analyses in the native brain space of the participants without spatial smoothing, as normalization and smoothing can induce spurious overlap between neighboring regions (Weiner and Grill-Spector, 2013).

We found that both OTS-words sub-regions prefer emojis over texts, which is surprising, as most prior studies have argued for a selectivity for text in OTS-words (Cohen et al., 2000; Cohen & Dehaene, 2004; Szwed et al., 2011). However, other prior studies have questioned this exclusive preference for text, showing that activations in OTS-words can be elicited by other visual stimuli as well (Price and Devlin, 2003; Ben-Shachar et al., 2007; Mei et al., 2010). Here we identified the OTS-words subregions in the same fashion as previous studies (Stigliani et al., 2015; Rosenke et al., 2021; Keller et al., 2022) but in addition introduced emojis as a new “readable” visual stimulus. This allowed us to disentangle task and stimulus preferences, which is not possible if the only readable stimulus in the experiment is text. Hence, the difference between our findings and the prior literature may be explained by i) in our study text was compared to another readable stimulus, ii) we introduced a task that required participants to read all stimuli in the experiment, and iii) we presented emojis, with which the participants have prior experience and which relate to archaic forms of writing. We discuss each of these differences separately in the paragraphs below. The observed preference for emojis over text in mOTS-words and pOTS-words provides evidence that these regions are not solely selective for text but rather are flexibly recruited for processing any kind of readable stimulus.

The emoji stimuli presented in the current study differed from previous work in that they were readable, i.e., the two stimuli together formed meaningful English compound words. Similar to learning a new writing system, abstract stimuli can become “readable” when we learn to retrieve semantic meaning from previously insignificant visual features. For example, in a recent study subjects were trained to read “HouseFont”, an artificial writing system, in which English phonemes were presented as different houses (Martin et al., 2019). Strikingly, authors found higher activations in OTS-words for HouseFont than an artificial control script once training was completed. This aligns with our findings, suggesting that the OTS-words subregions are activated when semantic meaning needs to be assigned to an abstract stimulus. One important difference to our study is that Martin et al. used stimuli that initially were not related to the word that they represent, whereas our emoji stimuli were pictograms of the objects themselves. These differences raise the question which kind of stimuli can be turned into readable units. Future research could investigate if readability might also be evoked for stimuli such as photographs of objects or natural scenes. Another difference is that, in the study by Martin et al. each house was mapped onto a phoneme, which then needed to be combined into words. In our study, each emoji represented an entire word, and two emojis needed to be combined into a compound noun. Interestingly, analogous to these differences, across different scriptures, symbols can either represent individual letters (e.g., English), phonemes (e.g., Finnish), or words (e.g., Japanese kanji). This suggests that OTS-words might be surprisingly flexible regarding the mapping between symbols and language units.

Our findings highlight the importance of the performed task in driving responses in the OTS-words subregions. For example, while we did not find effects of stimulus or repetition in mOTS-words in experiment 2, we found both task and stimulus effects in experiment 3 where a reading task was introduced. This aligns with recent studies showing higher responses for reading tasks than other tasks in OTS-words (Grotheer et al., 2018, 2019; Qu et al., 2022). Moreover, the reading task employed in the current study may even have contributed to the surprising preference for emoji stimuli observed in OTS-words, as conjoining evidence from intracranial recordings (Nobre et al., 1998; Woolnough et al., 2021), EEG (Strijkers & Bertrand, 2015), and fMRI (Mano et al., 2013; Kay et al., 2015; White et al., 2023) suggest that the task relevance of a stimulus can affect visual processing in OTS-words. Importantly, as outlined in the Interactive Account hypothesis (Price and Devlin, 2011), these task effects may not originate in OTS-words, but rather result from top-down modulation from higher level regions in the language and/or attention networks, as supported by recent functional connectivity (White et al. 2023) and fMRI modelling analyses (Kay et al., 2015). An open question is what kind of task is needed to drive responses in OTS-words. Considering that OTS-words is also activated during object naming (Bookheimer et al., 1995; Moore and Price, 1999; Murtha et al., 1999) the question arises, if the observed preference for emojis is driven by reading or by silently naming the presented objects. However, recent work has indicated higher responses for reading than object naming in OTS-words (Quinn et al., 2017) and as our participants had to combine two emojis into a semantic unit, the reading task developed in the current study likely required a higher level of lexical processing that goes beyond object naming.

The observed preference for emoji stimuli in OTS-words may also be explained by the extensive experience our participants have with emojis. Previous studies showed higher activations in OTS-words for familiar compared to unfamiliar scripts (Szwed et al., 2011; Dehaene et al., 2015; Martin et al., 2019), highlighting the importance of training in driving OTS-words. Indeed, OTS-words is absent in illiterate adults (Dehaene et al., 2010, 2015; Braga et al., 2017) and proposed to emerge due to training during our childhood education (Ben-shachar et al., 2011; Dehaene-Lambertz et al., 2018; Nordt et al., 2021). As our participant stem from a demographic that likely encounters emojis daily (Prada et al., 2018), and as emojis convey critical social information (Kaye et al., 2017; Boutet et al., 2021) expertise for emojis might hence have contributed to the observed preference in OTS-words. Moreover, while most modern scripture systems are based on morpheme and phonographic signs, the earliest scripture systems, the Sumerian cuneiform and the Egyptian hieroglyphs, were iconographic (Trigger, 1998). They were composed of pictograms, expressing objects, actions and ideas, similar to modern day emojis. Emojis may hence actually represent a prior, more fundamental way of written communication (Alshenqeeti, 2016). Nonetheless, as we observed longer response times when participants read emojis compared to text, one might still argue that reading emojis leads to an increased cognitive load resulting in higher responses in OTS-words. More research is required to fully understand why OTS-words shows a preference for emojis compared to text, for example by evaluating if there are distinct populations of voxels within these regions that selectivity process one stimulus or the other. After all, the fact that we did not observe fMRI adaptation in the mixed condition in Exp 2 and that we could reliably decode the visual stimulus in both subregions across experiments may support this scenario.

We conclude that both OTS-words sub-regions contribute to fluent reading, and that they are flexibly recruited whenever semantic meaning needs to be assigned to readable visual stimuli. Our work further suggests that both stimulus features and task demands should be considered when investigating the OTS-words subregions.

## Acknowledgements

This research was supported by the National Eye Institute (R01EY033835), by the Deutsche Forschungsgemeinschaft (DFG, German Research Foundation – project number 222641018 – SFB/TRR 135 TP C10), as well as by “The Adaptive Mind ‘ ‘, funded by the Excellence Program of the Hessian Ministry of Higher Education, Science, Research and Art.

